# Localization and RNAi-driven inhibition of a *Brugia malayi* encoded Interleukin-5 Receptor Binding protein

**DOI:** 10.1101/2021.05.13.443627

**Authors:** Rojelio Mejia, Sasisekhar Bennuru, Yelena Oksov, Sara Lustigman, Gnanasekar Munirathinam, Ramaswamy Kalyanasundaram, Thomas B. Nutman

## Abstract

A molecule termed BmIL5Rbp (aka Bm8757) was identified from *Brugia malayi* filarial worms and found to competitively inhibit human IL-5 binding to its human receptor.

After the expression and purification of a recombinant BmIL5Rbp and generation of BmIL5Rbp-specific rabbit antibody, we localized the molecule on *B. malayi* worms through immunohistochemistry and immunoelectron microscopy. RNA interference was used to inhibit BmIL5Rbp mRNA and protein production. BmIL5Rbp was shown to localize to the cuticle of *Brugia malayi* and to be released in their excretory/secretory products. RNAi inhibited BmIL5Rbp mRNA production by 33% and reduced the surface protein expression by ~50% and suppressed the release of BmIL5Rbp in the excretory/secretory products. RNAi has been used successfully to knock down the mRNA and protein expression of BmIL5Rbp in the early larval stages of *B. malayi* and provided a proof-of-principle for the local inhibition of the human IL5 receptor. These findings provide evidence that a parasite encoded IL5R antagonist could be utilized therapeutically.

## Introduction

Filarial parasites are vector-borne nematodes that are responsible for >200 million infections, the most pathogenic of which are caused by *Wuchereria bancrofti, Brugia malayi* (two of the causative agents of lymphatic filariasis), *Onchocerca volvulus* (the causative agent of onchocerciasis or “river blindness”) and *Loa loa* (the causative agent of loiasis or “the Africa eyeworm”). Infection occurs when third-stage larvae (L3) are introduced into the skin, after which they migrate through the dermis into either deep or superficial lymphatics, where they continue their development. With the identification of innate lymphoid cells (ILC) (1) and tissue-resident eosinophils in the skin (2) and the fat associated subcutaneous tissue, and the fact that the filarial L3s can evade these crucial innate cell populations to establish infection suggests that there must be a common mechanism used by these skin-transiting parasites to establish infection in humans. This lack of response is believed to be due to the parasite’s evasive immune mechanisms, especially the third-stage larvae (3).

Several helminth-encoded cytokine and chemokine mimics have been identified, largely through *in silico* analyses of genomic information. These include migration inhibitory factor (4–6) and TGF-β (7–9). A number of additional studies have used other methods to identify parasite-encoded molecules that are either agonists or antagonists of host-derived cytokine. Indeed, we identified in 2004, using a *B. malayi* L3 phage display library (10) a parasite-produced molecule called BmIL5Rbp (subsequent genome annotation Bm8757).

To understand the precise localization of this molecule in the *B. malayi* L3 and explore the feasibility of using L3s that fail to express this molecule for *in vivo* assessment/functional capabilities in relevant animal models, we used immunostaining and confocal microscopy to demonstrate the location and expression of BmIL5Rb. Using the RNAi approach on adults and microfilariae of *B. malayi* (11–14), we demonstrate its utility in L3s so that its role could be elucidated at the host-parasite interface.

## Results

### BmIL5Rbp is located on the surface of the worm and in the excretory-secretory products

Using immunostaining to localize BmIL5Rbp on various stages of *B. malayi*, it can be seen that BmIL5Rbp was found to be expressed on the surface of the worm using electron microscopy (Figure 1) and laser confocal microscopy of the adult male, female, and L3 (Figure 2). As seen, BmIL5Rbp (red) is only demonstrated in the stacked images on the parasite surfaces. Interestingly, the protein is found dispersed throughout the worm’s cuticle. No BmIL5Rbp was seen internal to the cuticle even when using fragmented or transected worms (data not shown).

**Figure 1.**
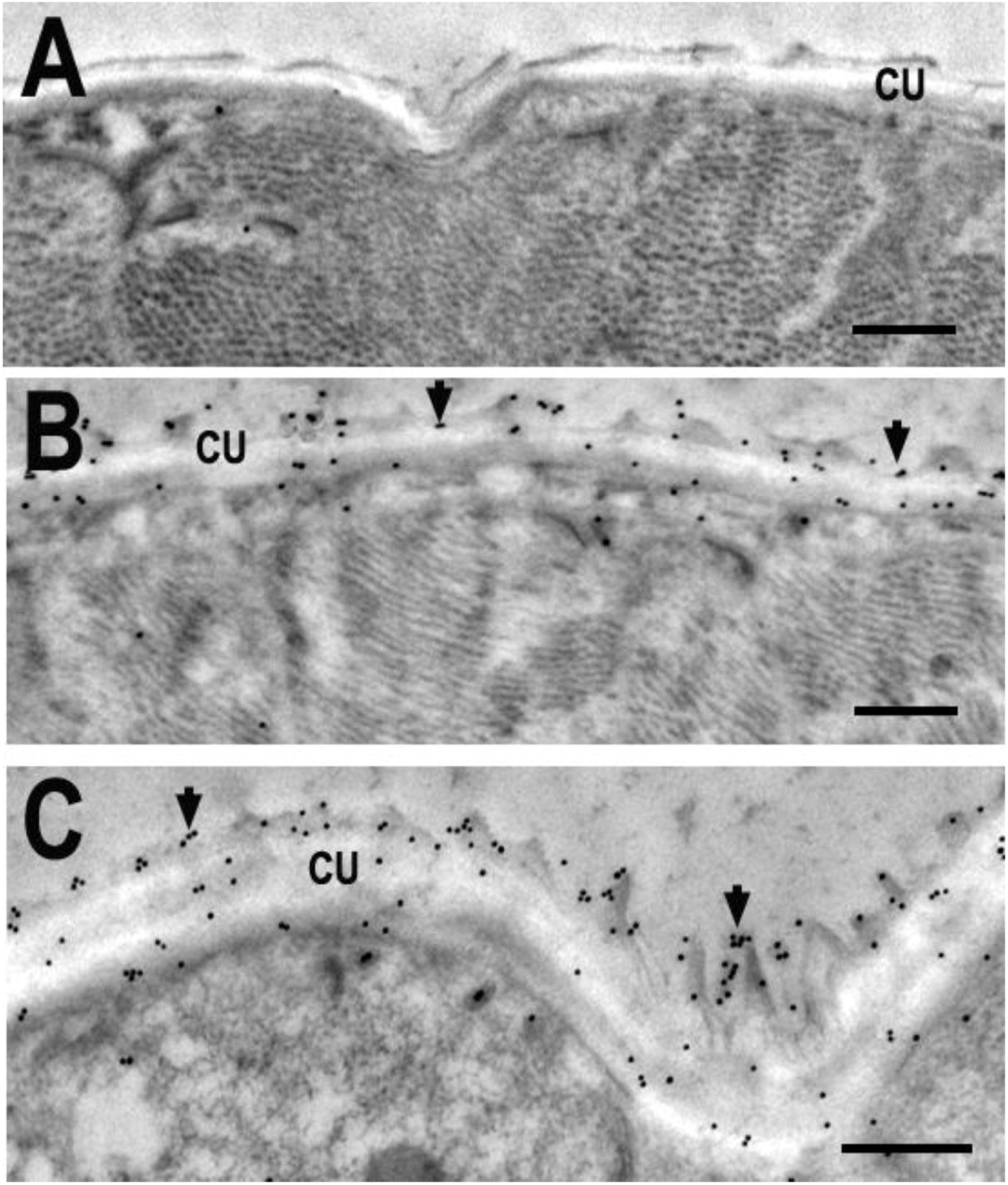
Evaluation of *Brugia malayi* L3 larvae sections under a transmission electron microscope showed that BmIL5Rbp is predominantly concentrated on the surface of the parasite (arrows), suggesting that BmIL5Rbp is expressed on the parasite’s surface. The micrograph (A) shows worms coated with preimmunized rabbit sera against BmIL5Rbp compared to the micrograph (B, C) using post-immunized rabbit sera identifying antibody localization on the worm’s cuticle. Micrograph (C) is at higher magnification with Bar 500 nm.

**Figure 2.**
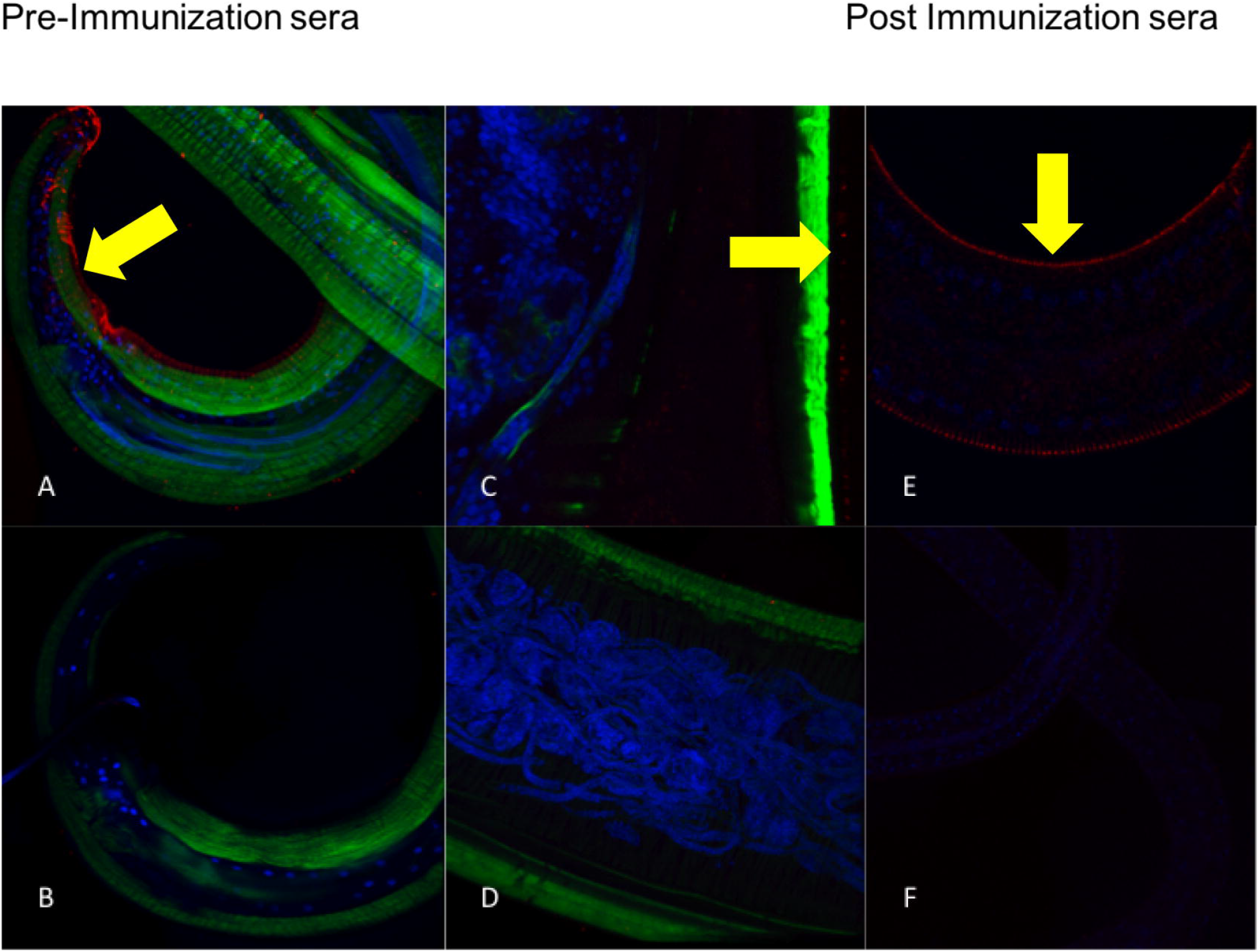
Immunohistochemical staining localization with post-immunization rabbit sera to BmIL5Rbp and laser confocal microscopy on *Brugia malayi*. Worms were stained for BmIL5Rbp (red), nuclei (blue), and actin (green). Adult male worms (A, B), adult female worms (C, D), and infectious stage L3 worms (E, F) all express BmIL5Rbp on the surface of the cuticle. No BmIL5Rbp was seen internally in the worms, even dissected specimens. Control worms (B, D, F) had minimal red staining with pre-immunized rabbit sera to BmIL5Rbp.

### Silencing BmIL5Rbp mRNA in L3 worms

To undertake the silencing of the BmIL5Rbp in L3s, we soaked L3 worms for two days in dsRNAi constructs for BmIL5Rbp or BmCPL (control) and examined the mRNA expression of the BmIL5Rbp. As shown in Figure 3A, BmIL5Rbp mRNA expression was diminished significantly compared to the BmCPL inhibited worms (33.1% vs. 4.89%, p = 0.0007) (Figure 3A). Approximately 85% of worms survived in both groups. Supernatants from live worms used in these RNAi cultured showed that the BmIL5Rbp protein content from these silenced worms showed levels of BmIL5Rbp (Figure 3B) that were 50% of the control worms (GM 205 pg/ml for the control siRNA construct and GM 105 pg for the siBmIL5Rbp group (P=0.034).

**Figure 3.**
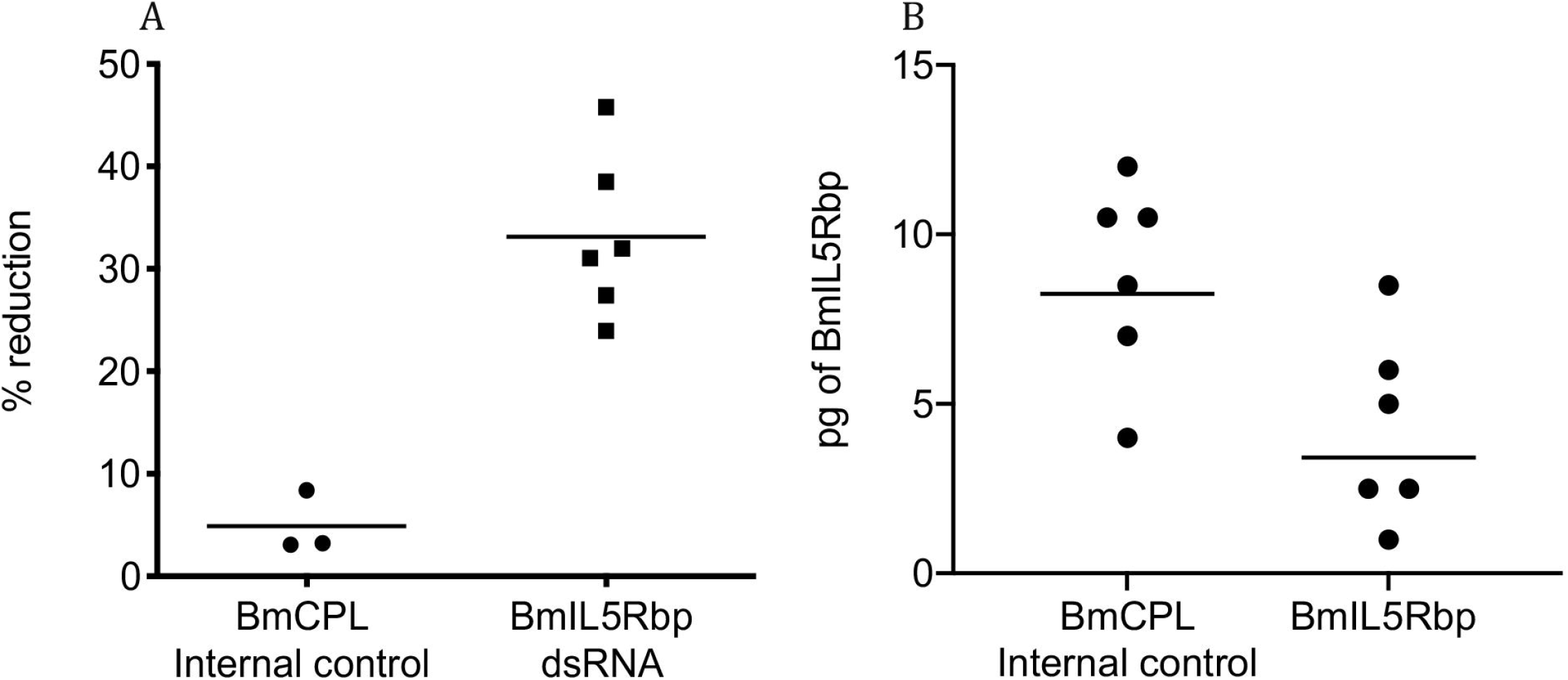
(A) BmIL5Rbp mRNA expression in live *Brugia malayi* L3 larvae is inhibited using RNAi. RNAi-specific BmIL5Rbp dsRNA had a significant percent reduction to the internal control BmCPL dsRNA (33.1% versus 4.9% respectively, p = 0.007). A total of 450 worms were processed with an 85% survival rate in both groups. (B) RNAi inhibits BmIL5Rbp protein expression in excretory/secretory products of live *Brugia malayi* L3 larvae compared to control worms (3.4 versus 8.2 pg/μL respectively, p = 0.034).

### Detection and quantification of surface and excretory/secretory BmIL5Rbp

There was a 54% decrease in surface protein expression between the two groups of worms, as shown by the confocal images in Figure 4A. Quantification of fluorescence at the surface of the worms (Imaris) showed a similar reduction induced by siBmIL5Rbp (2.125 Intensity Sum/Voxels compared to 1.146 Intensity Sum/Voxels (p = 0.0025).

**Figure 4.**
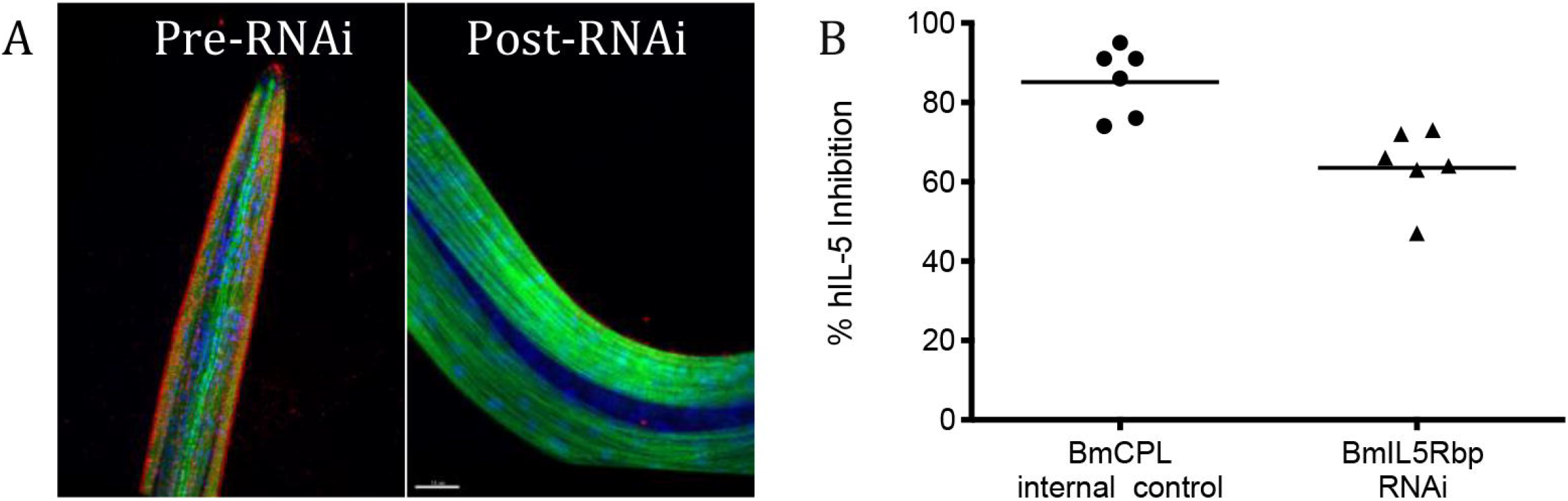
(A) RNAi can effectively reduce BmIL5Rbp on the surface of *Brugia malayi* L3 larvae. Intensity sum is the calculation of all red-stained BmIL5Rbp normalize to the total number of pixels (voxels) (Intensity Sum/Voxels) of all stained components of the worm. Pre-RNAi worms showed significant BmIL5Rbp staining to post-RNAi worms (2.125 Intensity Sum/Voxels compared to 1.146 Intensity Sum/Voxel respectively, p = 0.0025). (B) The supernatant (sups) from RNAi *Brugia malayi* L3 larvae contain less BmIL5Rbp in their excretory/secretory product compared to internal controls and thus had less inhibition of human IL-5 to its human receptor (63.5% versus 85.1% respectively, p = 0.005).

Human IL-5 exhibited decrease binding to human IL-5 receptors with increasing rBmIL5Rbp concentrations (Spearman r = −0.9977, p < 0.0001) (Supplementary Figure 1). RNAi worms excrete less BmIL5Rbp to inhibit binding of human IL5 to its receptor. Supernatant from worms (200 in each group) at two days of soaking showed a decrease in inhibition of human IL-5 for the BmIL5Rbp RNAi worms compared to the negative control worms by 23% (Figure 4B). Results are presented in percentage of inhibition to control worms.

## Discussion

Lymphatic filarial parasites are notorious for their ability to manipulate host immune responses. In the present study, we show that one such mechanism includes interfering with the function of host-derived IL-5. Here we characterize and localize the BmIL5Rbp produced by the third stage infective larvae of *B. malayi*, a secreted molecule that can bind to soluble human IL-5 receptor alpha and block the binding of human IL-5 to this receptor. Thus, BmIL5Rbp, a protein produced by *B. malayi*, appears to be a novel host-immunomodulatory molecule, acting as an IL5 antagonist.

The ability of a nematode-derived protein to alter the human host immune response is a unique evolutionary response that likely helps the worm to establish a foothold. We describe a nematode-produced protein found on the worm’s surface (Figures 1, 2) and in its excretory/secretory product (Figures 3, 4).

*B. malayi*, along with all of the pathogenic filariae, induce a robust eosinophil response, particularly in the acute phase of infection (19, 20) related to the tissue migration L3 and L4 larvae of filarial nematodes (21–23). Our data suggest that these parasites secrete a molecule that antagonizes the activity of human IL-5, a cytokine found both early in infection and within days of anthelmintic chemotherapy (24–26).

The localization of BmIL5Rbp to the worm’s surface appears to be directly on (and within) the cuticle. Its location gives it accessibility to directly tissue-dwelling host cells bearing the IL5R (eosinophils, basophils, mast cells). It suggests that there may be a physiologic role for this molecule at the host tissue level. Since eosinophil worm killing is contact or localized dependent action (27), then the immediate areas, including the tissue, around the worm’s surface will have higher concentrations of BmIL5Rbp, leading possibly to a loss of eosinophil survival at the site of inflammation/tissue damage.

The use of RNAi to selectively inhibit BmIL5Rbp helps to provide proof of principle that the worm produces a protein to inhibit the binding of human IL5 to its receptor. The RNAi-treated worms produced specifically less BmIL5Rbp that had less inhibition of human IL-5 to its human IL-5 receptor *in vitro*. With the advent of CRISPR-CAS9 in genomic splicing (28), the ability to knock out this gene now is a possibility (29–31). However, for the filarial nematodes, RNAi remains the most useful method of gene silencing (11, 12, 14, 17, 32–34).

## Conclusion

While this study shows that it is feasible to silence the BmIL5Rb expression, *in vivo* experiments will be required to demonstrate the importance of this molecule (13). Our study provides a framework for utilizing these parasite-encoded cytokine antagonists for mechanistic and/or host-directed therapy studies that should, in turn, provide new insights into specific host-parasite interactions.

## Methods

### Expression and purification of rBmIL5Rbp

The entire coding sequence (Supplementary Table 1) of the IL5Rbp was amplified and inserted into entry clones containing an upstream Tev protease cleavage site (ENLYFQG). Both entry clones were completely sequenced to verify that they contained the correct sequence. Expression clones were transferred to six different T7 promoter-based Destination vectors for *E. coli* expression. Among the six, one termed 2110-X1-566 contained His6-MBP-tev-I5RBP that was found to produce the most abundant soluble protein when transformed into *E. coli* Rosetta (DE3). The soluble fraction of the MBP-BmIL5 was bound to the affinity column of the immobilized metal-ion affinity chromatography (IMAC) and eluted at 200 mM imidazole (~200 mM). The eluted material was incubated overnight with 5% TEV protease at 16° C. The contaminating components of the protease reaction were removed in a subsequent IMAC step. The final protein was ~ 95% pure as confirmed by SDS-PAGE analysis and was stored at 1mg/ml in PBS, pH 7.5 at −80° C until used. Recombinant BmIL5 (rBmIL5Rbp) was also expressed in Sf9 and Hi5 cells by infection with baculoviruses. 2 x 10^6^ insect cells from 48 post infection Hi5 cells incubated at 21°C were resuspended in lysis buffer (20 mM Na phosphate pH 7.5, 500 mM NaCl, 5% glycerol, Complete protease inhibitor, Roche Diagnostics, Rotkreuz, Switzerland) and sonicated until cells were > 99% lysed as determined by microscopic examination. Soluble and insoluble fractions were separated by centrifugation. Baculovirus-infected Hi5 cell paste was resuspended with extraction buffer (20 mM HEPES buffer, pH7.3, 500 mM NaCl, 45 mM imidazole, 5 mM MgCl_2_, 10% glycerol, 2 mm ßME, complete protease inhibitor [Roche] at 2 tablets/50 ml of lysate) to a volume in milliliters equal to four times the wet weight of the pellet in grams. The sample was sonicated to lyse the cells (verified by microscopic examination), treated with 1.0 U benzonase/ml for 20 min on ice, clarified by centrifugation at 111,000 x g for 50 min, filtered (0.45 μm PES membrane), diluted five-fold with extraction buffer and applied to a 5 ml HisTrap column (GE Healthcare, Chicago, IL) equilibrated with extraction buffer. The column was washed to baseline, proteins eluted over a 20-column volume (CV) gradient to 400 mM imidazole and fractions analyzed by SDS-PAGE. The pool created from the IMAC fractions was dialyzed against binding buffer without additional imidazole and treated with TEV protease (1% v/v with a lab stock) at RT for 4 hours and then shifted to 16 °C overnight and analyzed by SDS-PAGE. An additional IMAC step similar to the initial IMAC was used to purify the target protein away from the contaminants of the protease digest. Fractions were analyzed by SDS-PAGE.

### Electron microscopy and immunogold staining

The native BmIL5Rbp was identified in thin sections of *B. malayi* L3 by immuno-electron microscopy. The larvae were fixed with 4% paraformaldehyde/0.15% glutaraldehyde in 0.1 m sodium cacodylate buffer for 1 hour and embedded in LR White medium as previously described (15). Thin sections (70 nm) were then blocked with 2% BSA and incubated with primary antibody (rabbit anti-rBmIL5) or control serum (pre-immunization bleed of the same rabbit) overnight at 4°C. Following washes in PBS containing 0.1% BSA and 0.01% Tween-20, grids were incubated with gold-labeled goat anti-mouse IgG (Fc) at 1:20 dilution (Amersham Pharmacia Biotech, Little Chalfont, United Kingdom) for 1 hour at room temperature, washed, counterstained in 5% uranyl acetate solution and observed using a Philips 410 electron microscope (Figure 1).

### Immunostaining of L3 worms

L3 worms were fixed with a final concentration of 2% paraformaldehyde overnight at 4°C. Worms were washed three times with blocking buffer: 1% bovine serum albumin, 0.5% Triton X-100, 0.1% sodium azide, phosphate-buffered saline (PBS). The worms were soaked in 10 mMol sodium citrate (in blocking buffer) at 70°C for 60 minutes, washed three times in PBS, and soaked in blocking buffer overnight at 4°C. The worms were washed three times in PBS and then soaked in rabbit anti-BmIL5Rbp (1:10 in blocking buffer) for three days at 4°C. After washing three times in blocking buffer for 30 minutes, the worms were incubated with 1:500 dilution of goat anti-rabbit IgG Alexa Fluor 594 (Thermo Fisher Scientific, Waltham, MA) in blocking buffer for three days at 4°C. Phalloidin Alexa Fluor 488 (Thermo Fisher Scientific) in methanol (200 units/mL) was added and incubated overnight at 4°C. The worms were then washed three times with moderate shaking at 4°C for three hours each. Individual worms were collected and set with a mounting solution (Hardset VectaShield Vector H-1400, Vector Laboratories, Burlingame, CA) overnight a 4°C. All worms were visualized using a confocal microscope, Leica SP5 X-WLL (Mannheim, Germany). Dapi stain was detected at 460 nm, Phalloidin at 520 nm, Alexa Fluor 594 at 620 nm with excitation at 595 nm at 15% laser power, a smart gain of 838, offset −0.3%.

### Synthesis of siRNA constructs for BmIL5Rbp

Primers and probes for the gene sequence of BmIL5Rbp were designed using siRNA Target Designer-Version 1.6, Promega (Madison, WI) that used proprietary algorithms to select the optimal primer/probe sequences.

Forward 5’AAA AAG GGA ATT AGT TGT TG3’

Reverse 5’ATA CTA ACA AAC TTC TGC AAA TT3’

Plasmid DNA for BmIL5Rbp was generated and transformed to recombinant protein using TOP 10 *E. coli* competent cells (AMP resistant) (Thermo Fisher Scientific, Waltham, Ma). RNA PCR was performed using T7, T3 primers. Single-stranded RNA was annealed per-protocol (Agilent Technologies, Santa Clara, CA) with a final product of double-stranded RNA (dsRNA). DsRNA was concentrated using microcentrifugation filters and the final product was resuspended in PBS.

### RNAi of *Brugia malayi* L3 worms

The L3 worms were cultured, as described previously (16). Double-stranded BmIL5Rbp (dsBmIL5Rbp) RNA was introduced to L3 worms by soaking, in which 20 L3 worms were soaked in 100 μL of dsBmIL5Rbp (500 μg/mL) in L3 media (16) for two days at 37°C. The media was removed, and 100 μL TRIzol^®^ (Thermo Fisher Scientific) was added to the L3’s and quickly frozen. The L3 larvae went through three freeze-thaw cycles and homogenated for five minutes at room temperature with Qiagen shredders (Qiagen, Venlo, Netherlands), washing with 100 μl TRIzol^®^ (Thermo Fisher Scientific) three times. The macerated larvae were centrifuged at 12,000 g for one minute at 4°C between each wash. A total of 20 μL chloroform was then added to the shredded larvae and shaken vigorously for 15 seconds. The mixture was then centrifuged for 15 minutes at 12,000 g at 4°C. The aqueous phase was retained, and equal volumes of isopropanol were added. The aqueous phase mixture was centrifuged at 16,000 g for 15 minutes at 4°C. The pellet was washed with 70% ethanol then centrifuged at 16,000 g for 5 minutes at 4°C. The pellet was air-dried for 20 minutes, then re-suspended with 10 μL TE buffer (Thermo Fisher Scientific). Reverse transcription PCR was performed using standard techniques to the final volume of 100 μL. An internal control siRNA for *Brugia malayi* cathepsin L intron (BmCPL) (17) was created using the above protocol and was used to show specific inhibition of BmIL5Rbp using a SYBR Green real-time PCR (Applied Biosystems) per manufacturer specifications.

Primers for BmCPL (14).

Forward: 5’ GAC AAA GAT TAC AAA CAG GGC3’
Reverse: 5’ TGA TTG GGC AGT CGA AGT C3’

### Real-time PCR quantification of BmIL5Rbp

Real-time PCR with final volumes of 20 μL, 1 μL of the probe (FAM-5’AAG CAA GCA TTG ATT TCT3’-MGBNFQ), 2 μL of template, 10 μL Taqman^™^ fast mix (Applied Biosystems, Foster City, CA), forward/reverse primers 1 μL each, 5 μl of dH_2_O. Primers for BmIL5Rbp were selected upstream of the siRNA selection using the following primers:

Forward: 5’AAA ATG ATG GAA GCA GCA GAA ACT G3’
Reverse: 5’TAC AGG AAT ACA TCA TGC TCA CAA GT3’

All reactions were performed on an ABI 7900HT Fast Real-Time PCR System (Applied Biosystems). All primers and probes were purchased from Applied Biosystems. The PCR conditions were 95°C for 1 min 1 x cycle; 95°C for 25 s, 55°C for 20 s, 72°C for 1 min 35x cycles; 72°C for 10 min 1x cycle.

For standard comparison, housekeeping control Bm-tub primers were used with SYBR Green real-time PCR (Applied Biosystems) per manufacturer specifications (14).

Primers for Bm-tub (17)

Forward: 5’ATA TGT GCC ACG AGC AGT C3’
Reverse: 5’CGG ATA CTC CTC ACG AAT TT3’

### Penetration of dsBmIL5Rbp in live *Brugia malayi*

DsBmIL5Rbp was Cy3 labeled using the CyDye^™^ Post-Labelling Reactive Dye Pack (GE, Piscataway, NJ) per manufacturer’s instructions. A total of 5 μL dsBmIL5Rbp (2000 μg/mL) was added to 100 mMol sodium bicarbonate and mixed with Cy3 in 20 μL DMSO stirred in the dark for 90 minutes at room temperature. Then 15 μL of 4M hydroxylamine was added and mixed for 15 minutes in the dark at room temperature. Next, the mixture was precipitated with 5 μL of 5M sodium chloride for 60 minutes at 20°C. The solution was centrifuged at 16,000 g for 15 minutes at 4°C. Then to the supernatant, 175 μL of 70% cold ethanol was added, then centrifuged 16,000 g for 5 minutes at 4°C. Next, the supernatant was removed, and the pellet air-dried for 10 minutes in the dark and solubilized in 100 μL in L3 media (16). A total of 20 live L3 worms were soaked in the dsBmIL5Rbp media overnight. The worms were then fixed and mounted at the above methods. Also, 20 other worms were used in non-CY3 L3 media as a control group

### Quantifying surface BmIL5Rbp on L3 worms using confocal microscopy

Using the above RNAi methods, approximately 60 worms were exposed to BmIL5Rbp RNAi constructs. These were mounted along with appropriate control worms. Using a laser confocal microscope, capturing several 0.5mm stack images were obtained of entire worms. The intensity was calculated by using sum/voxels in Imaris (South Windsor, CT). Voxels are the total number of pixels at a specific wavelength.

### Quantification of excretory/secretory BmIL5Rbp in L3 worms

BmIL5Rbp was measured using an immune dot blot. Briefly, nitrocellulose paper (Schleicher and Schuell (Keene, NH). was trimmed to fit a 96-well manifold. The paper was activated in methanol for three minutes and then washed for 1 min in distilled water. The paper was dried, and 50 μL of rBmIL5Rbp (baculovirus derived) in various dilutions, duplicates were placed in wells with 50 μL of supernatants from RNAi and control worms. The plate was incubated at 37°C for one hour. The plate was dried, and the membrane was placed in a blocking buffer (5% milk, 5% rabbit serum, 5% goat serum) overnight at 4°C with gently tilting action. The membranes were incubated at room temperature in primary antibody Rabbit anti-BmIL5Rbp 1:100 dilution in 5% milk solution. The membrane was washed three times with 0.5% Tween 20 (Sigma-Aldrich, St. Louis, MO) in PBS solution for 5 min each cycle. Then a secondary antibody Goat anti-Rabbit Horseradish peroxidase (HRP) (Thermo Fisher Scientific) 1:1000 dilution in 5% milk was added for a 1-hour incubation at room temperature. The membrane was washed three times with 0.5% Tween and PBS solution for 5 minutes each cycle. For the ELISA developing step, a SuperSignal West Pico Chemiluminescent Substrate kit (Thermo Fisher Scientific) was used on the membrane as per the manufacturer’s instructions. Luminescence was detected using Kodak X-OMAT film (Rochester, NY) and exposed for 10 minutes, then quantitated using BmIL5Rbp standard curves.

### Measurement of decrease inhibition of human IL5 to its receptor *in vitro* using RNAi worms

A total of 50 μL/well of recombinant human IL5-Receptor alpha (R&D Systems, Minneapolis, MN) final concentration of (4 μg/mL) was plated using coating buffer (18) on Immulon 4 plates (Thermo Fischer Scientific) overnight at 4°C. The plates were then washed with (0.2% Tween 20 in PBS) 6 times. A total of 100 μL of ELISA blocking buffer (0.05 Tween 20, 5% bovine serum albumin, PBS) was added for 1 hour at 37°C, with subsequent washing as above. Afterward, 50 μL of E/S product from 200 worms in each group (BmCPL Internal control and BmIL5Rbp RNAi) was added and incubated for 1 hour at 37°C. The plates were washed again as above. Subsequently, 25 μL of biotinylated recombinant human IL5 (rhIL5) protein (50 μg/mL) (R&D Systems) was added at 1:50 in PBS. Biotinylated rhIL5 was produced using the Biotin-X-NHS system (Merck, Darmstadt, Germany). A total of 25 μL of rhIL5 (100 μg/mL) was immediately added for a final concentration of 50 μg/mL and incubated at 37°C for one hour. The plate was washed as above, and 50 μL of streptavidin (Thermo Fisher Scientific) was added and incubated at 37°C for one hour. The plate was washed for a final time as above. The plate was developed with pNPP phosphatase substrate (Sigma, St. Louis, MO) per the manufacturer’s instructions for 30 min at room temperature. Plates were read at 405 nm in an ELISA reader (SpectraMax Plus; Molecular Devices, Sunnyvale, CA). Similar methods were used with increasing concentrations of rBmIL5Rbp inhibiting hIL-5 to human IL-5 receptor (Supplementary Figure 1). In determining the optimal concentration for Biotinylated rhIL5, a separate set of experiments was done with 2-fold dilutions using the above methods. Results showed that a 1:100 final concentration of biotinylated rhIL5 was the optimal concentration (data not shown).

### Statistics and calculations

All statistical analysis was performed using Prism version 5.0d (GraphPad, La Jolla, CA). Comparisons were analyzed using the Mann-Whitney test, and P values < 0.05 were considered significant. RNAi results were shown as the percent reduction compared to a standard housekeeping control gene (Bm-tub) as previously described (17). In summary, the control group value was set as the maximum amount with the RNAi-treated worms reported as a relative reduction compared to the internal control group (BmCPL). Calculation of intensity sum/voxels was completed with Imaris Software (Zurich, Switzerland).

## Notes

### Competing Interest Statement

The authors have declared no competing interest.

