## Supplementary for "Localization and RNAi-driven inhibition of a *Brugia malayi* encoded Interleukin-5 Receptor Binding protein"

Supplementary data.

Table 2. BmIL5Rbp DNA sequence.

```
1    ATGAAAATGA TGGAAGCAGC AGAAACTGCT ATGAAAAAGC AAGCATTGAT TTCTACAATG
 61   AATGAGCAAC AGATACAGGA ATACATCATG CTCACAAGTC ATATAGAACA AGAACTAATA
121  AAAGAAAATC AAGAAAAATT CGCTGCAAAA AGGGAATTAG TTGTTGCGAA AGGGCAGCGA
181  AAAAATAAAG AAGAATATGA ATTATTGGCC AAAATGATTG AAAAAGTGCC GTCTAGATCA
241  GAAAGCACCA GGAAACTAGA CCTAATCAAG AAGGAACTTG AGGAACTACA TGAGAAGCAA
301  CGAAATTTAG AAACAAAGCT AGCGGAACGT CGCAATCATC TTCATTCTCT CAATATCATA
 361  CTAACAAACT TCTGCAAATT TTTGAAAGAC GAAGAGAACG GTGAAATAAT GAATAATGAT
421  GATGCTGTGG ACGGCACAGG GACAGATGGT AAAAAGGAGA AAGCTTGCGG CCGCACTCGA
 481  GTAACTAGTT AA
```

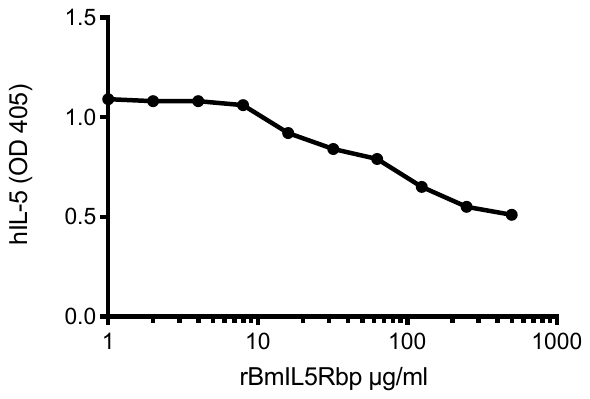

Supplementary Figure 1. Increasing concentrations of recombinant BmIL5Rbp inhibited human IL-5 to human IL-5 receptors using an inhibition ELISA (Spearman r = -0.9977, p < 0.0001).
